# Natively entangled proteins are linked to human disease and pathogenic mutations likely due to a greater misfolding propensity

**DOI:** 10.64898/2026.04.15.718104

**Authors:** Maria F. Anglero Mendez, Ian Sitarik, Quyen V. Vu, Prabhat Totoo, James D. Stephenson, Hyebin Song, Edward P. O’Brien

## Abstract

A recently discovered class of protein misfolding involving native entanglements could be a widespread mechanism by which loss-of-function diseases arise. Here, we test that hypothesis by examining if there is any statistical association between proteins predisposed to misfold in this way and a database of gene-disease relationships. We find that globular proteins containing non-covalent lasso entanglements (NCLEs) in their native structure, which are more prone to misfolding, are 61% more likely to be associated with disease, 68% more likely to harbor pathogenic missense mutations, and their misfolding-prone entangled regions are 64% more likely to harbor pathogenic missense mutations. Protein refolding simulations indicate that these disease associated, natively entangled proteins are 2.5-times more likely to misfold than comparable non-disease proteins that lack native NCLEs. These results indicate that native entanglement misfolding, especially in the presence of missense mutations, have the potential to contribute to a wide variety of diseases. More broadly, these findings open an entirely new space of therapeutic targets in which drugs are designed to avoid these misfolded states and increase the amount of folded, functional protein.

## Introduction

In many well-known diseases proteins gain new abilities and functions^1,2^. In Alzheimer’s and sickle cell anemia, proteins acquire the ability to undergo uncontrolled aggregation^3,4^. Most diseases, however, are not caused by new aberrant behavior, but instead caused by proteins losing their ability to function normally^2^. There are a number of ways this can happen. Mutations can cause a protein to misfold and malfunction, be excessively degraded and reduce the total amount of functional protein, or destroy the key residues essential to their function^5–7^. Proteins can also fail to be fully synthesized resulting in incomplete sequences with no function^8,9^. Regardless of the mechanism, by losing enough functional protein, the subcellular and cellular processes they take part in can become impaired, and in some cases lead to disease. Mutations in the cystic fibrosis protein, for example, can cause it to misfold^7,10,11^ and result in a lower final amount of functional protein. This decreases ion flux in lung tissues and leads to a buildup of mucus. In the enzyme phenylalanine hydroxylase, certain mutations destabilize the protein making it harder to fold and reducing its ability to convert one amino acid to another^12,13^ resulting in neurotransmitter and hormonal deficits. Indeed, a large proportion of pathogenic mutations are associated with disease because they destabilize the native structure of the protein leading it to not properly fold and function^14^.

Recent studies indicate there is a previously unrecognized, widespread class of protein misfolding that can result in loss of function in proteins^15–20^. This type of misfolding, therefore, could be a common cause of these loss-of-function diseases. This hypothesis predicts that there should be a statistical association between diseases, pathogenic mutations, and proteins that are most susceptible to misfold in this way. In this study, we test if such associations exist.

The new class of misfolding involves proteins containing non-covalent lasso entanglements (NCLEs), which are geometric motifs consisting of a protein backbone loop closed by non-covalent interactions between residues and pierced by N- or C-terminal backbone segments^16–18^ (Fig. 1). If a protein’s native state contains one or more of these motifs it is twice as likely to misfold than similar globular proteins that lack them^16^. Between 49% and 68% of human globular proteins contain these NCLEs in their native structures^15^. (The numbers vary depending on whether the counting is done on the dataset of X-ray crystal structures or the larger set of AI-predicted AlphaFold2 structures.) Simulations indicate that misfolding of NCLEs occurs via its loop prematurely closing before the N- or C-terminal segments are properly positioned within the loop, creating steric and entropic barriers to those segments piercing the loop^16^. This mechanism is called the failure-to-form mechanism, because the NCLE fails to fold while the rest of the protein properly does so. Another less common mechanism was also observed in which an entanglement forms that is not present in the native state, leading to the gain of a non-native entanglement^16,17^. In both mechanisms, the resulting misfolded structure is often ‘off-pathway’ meaning it cannot directly convert into the native state but instead must unfold for the protein to have the opportunity to attain its native structure. This can cause these misfolded states to be long lived, kinetically trapped states, which are metastable yet persist from hours to days or longer^21,22^.

**Fig. 1:**
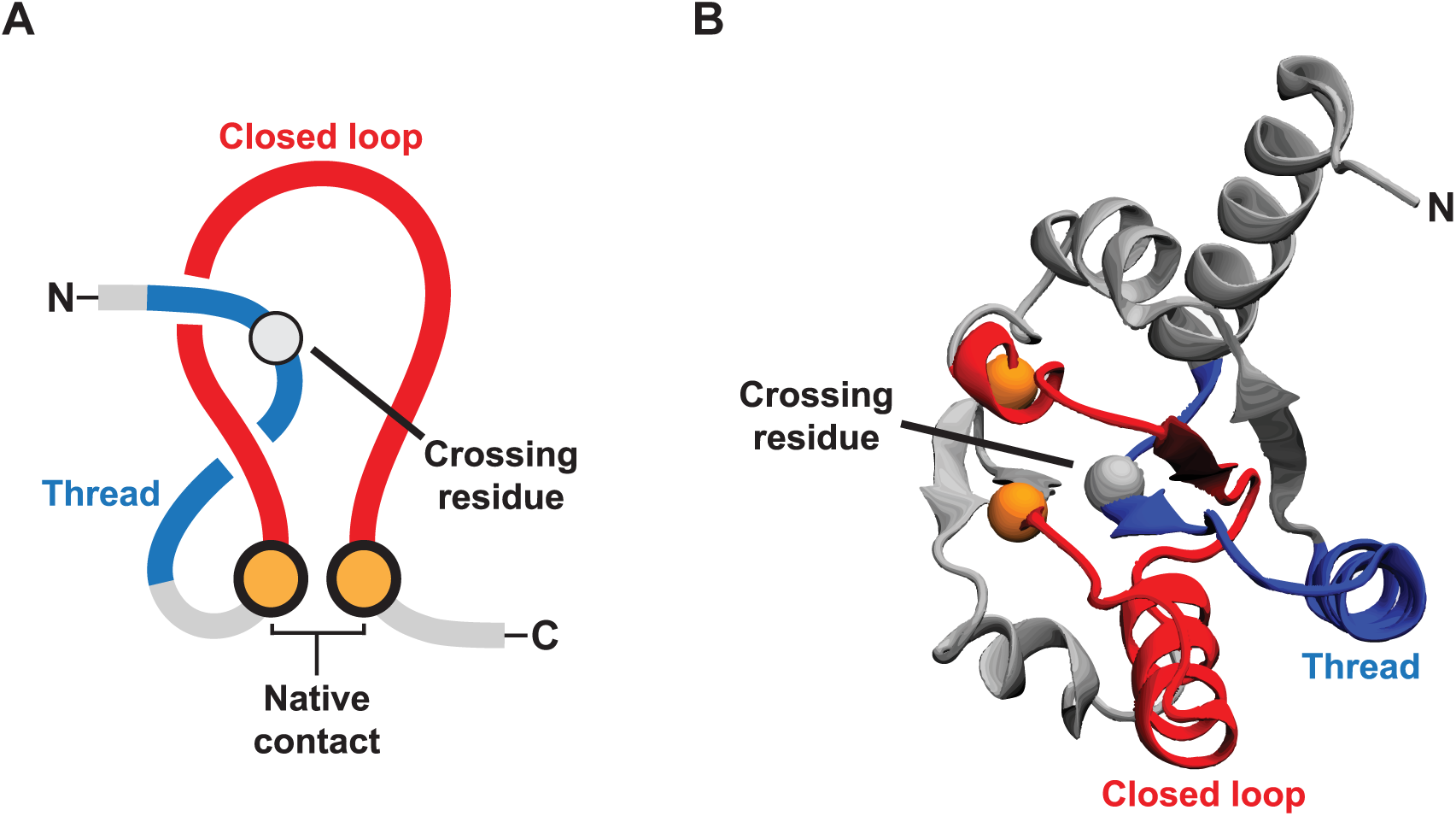
Illustration of the essential elements of a non-covalent lasso entanglement (NCLE). **a,** An NCLE consists of a loop (red) closed by a non-covalent contact (gold; residues with heavy atoms within 4.5 Å) and threaded by a segment outside the loop (blue)^24^. The crossing residue is the residue on the threading segment that pierces the loop plane. **b,** Example in the crystal structure of uL10 (P0A7J3; PDB 6XZ7, chain H), where a loop (red) closed by native contact between residues L72 and A111 (gold) is threaded by an N-terminal segment (blue), with a crossing at V27.

Some of these misfolded states can bypass the refolding action of chaperones^16,19^. Simulations indicate that proteins that exhibit this behavior do so by native NCLE’s misfolding into structures that are structurally similar to the native state^16,19^. This allows them to bypass chaperones to a similar extent as proteins adopting the native ensemble. These observations suggest a molecular mechanism – adoption of near-native misfolded states - by which such states can be long lived inside cells^23^.

Here, we examine if proteins with native entanglements – which are more likely to misfold^24^ – are statistically associated with (*i*) genes known to be associated with disease, (*ii*) pathogenic mutations, and (*iii*) particular disease classes. To gain insight into potential misfolding mechanisms we simulate the refolding of some fifty disease- and non-disease-causing proteins. Finally, we provide a list of proteins that, based on heuristics, are most likely to cause disease through NCLE misfolding.

## Results

### Entangled proteins are 61% more likely to be associated with disease

Globular proteins with native NCLEs are twice as likely to misfold as compared to those without these structural motifs^24^. We hypothesized that such misfolding could contribute to loss-of-function diseases. This hypothesis predicts that proteins containing native NCLES will be positively associated with genes known to be connected to disease. To test this prediction, we calculated both the adjusted odds ratio (Eq. S8), which controls for protein length and gene essentiality as confounding factors, and the crude odds ratio (Eq. S9). (The odds ratio quantifies the odds of disease associated proteins occurring in natively entangled proteins relative to the odds natively non-entangled proteins will exhibit disease association.)

Across our AlphaFold2^25^ structural dataset, 43.9% of proteins are associated with disease regardless of their native NCLE status. Of the proteins containing NCLEs, 49.5% are associated with disease, compared to 32.8% among those without a native NCLE. The crude odds ratio of association between the presence of one or more native NCLEs and disease is 2.01 (95% confidence interval (CI) = [1.86, 2.17], p-value= 3.66 × 10^−73^, Fisher’s exact test, Table 1), indicating a positive association. The adjusted odds ratio is 1.61 (95% CI = [1.48, 1.75]; p-value = 1.35 × 10^−30^, Wald test from logistic regression, Table S1). We conclude that proteins with native entanglements are more likely to be disease associated compared to those without these geometric motifs, increasing the odds by 61% (=(adjusted OR-1)*100%).

**Table 1.**
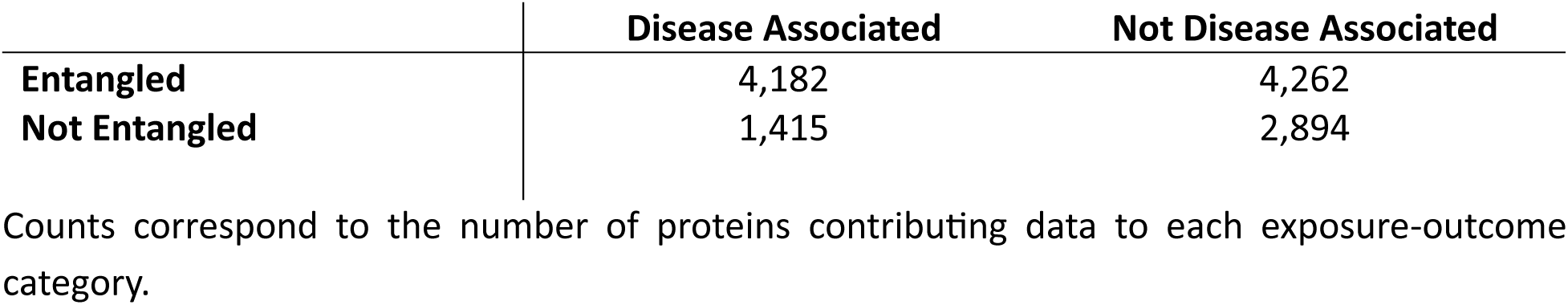
Distribution of disease association by entanglement status of the protein’s native structure.

### Entangled proteins are associated with particular disease classes

We next asked if this association varies between disease classes. A disease class is a group of diseases with shared pathological or biological characteristics. To address this, we analyzed the same 5,597 proteins associated with at least one disease and categorized them into the 23 disease classes defined in DisGeNET^26^. These classes group related diseases based on the MeSH disease categories under the “C - Diseases” and “F - Diseases” branches of the Medical Subject Headings (MeSH) thesaurus^27^. We then compared proteins in a particular disease class against all other proteins from DisGeNET and Uniprot^28^ that were not classified as disease associated for that specific class. We find that 18 of these disease classes exhibit positive associations with the presence of native entanglements (Eq. S10, Table S2, Fig. 2). The largest associations are for Behavior and Behavior Mechanisms (F01), Nutritional and Metabolic Diseases (C18), and Digestive System Diseases (C06) with adjusted odds ratio’s, respectively, of 1.74, 1.72, and 1.65. In the case of Behavior and Behavior Mechanisms (F01), for example, 10.7% (=597/5,597*100%) of entangled proteins are associated with this disease class, and only 3.4% (=145/4,309*100%) of non-entangled proteins are. Thus, entangled proteins exhibit widespread, significant positive associations with most disease classes.

**Fig. 2:**
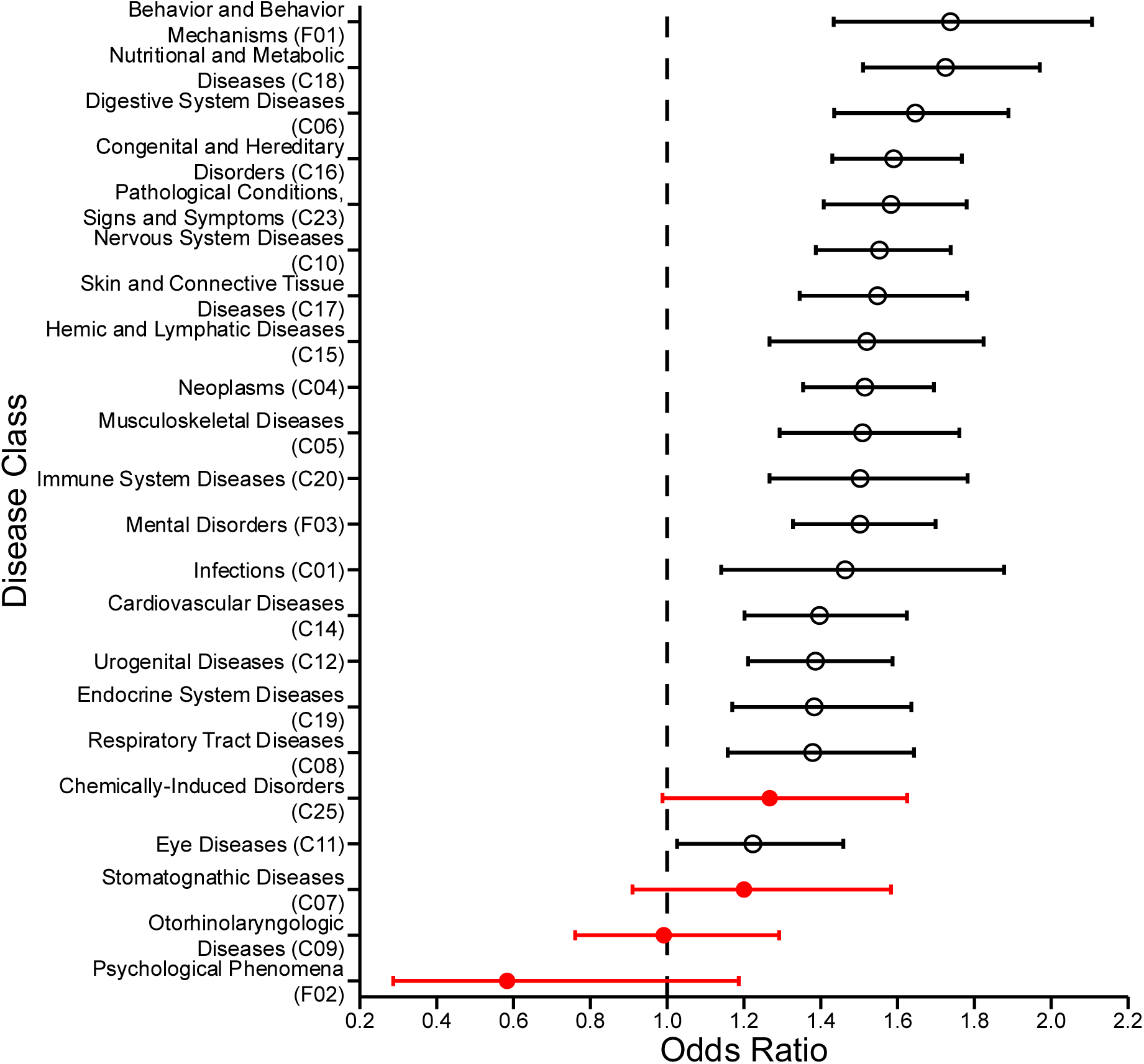
Odds ratios of native entanglement status and disease association by MeSH disease class. Odds ratios and 95% confidence intervals for the association between protein entanglement and disease status across 22 MeSH disease classes (Table S4). Points represent adjusted odds ratios from logistic regression models controlling for protein length and essentiality. Error bars indicate 95% confidence intervals. The dashed vertical line denotes an odds ratio of 1. Black open circles indicate statistically significant associations, and red filled circles indicate non-significant associations. Note: Disorders of Environmental Origin (C21), is not shown because its confidence interval extended to infinity (i.e., was undefined). Congenital, Hereditary, and Neonatal Diseases and Abnormalities (C16) was abbreviated to Congenital and Hereditary Disorders (C16) for display purposes.

### Structural properties of entanglements are associated with disease

Structural properties of proteins have been shown to correlate with the likelihood of protein misfolding^29^. Therefore, we hypothesized that the structural properties of native entanglements (Table S3) can affect the likelihood of misfolding – for example, more complex entanglements might be more likely to misfold - and therefore the likelihood of disease. To test this hypothesis, we calculated 17 structural metrics for each entangled protein in our dataset that describe the geometry, depth, and heuristic correlates of stability of native entanglements (Table S4). We then performed logistic regression to calculate the odds ratio that each metric is associated with disease, controlling for protein length and essentiality (Eq. S11). We indeed observe a number of entanglement features show positive associations with disease (Fig. 3), with the largest odds being 1.53 for the property *d*_P_(C), which describes the depth of the C-terminal crossings along the primary structure from the C-terminus (in residues), normalized by the length of the protein.

**Fig. 3:**
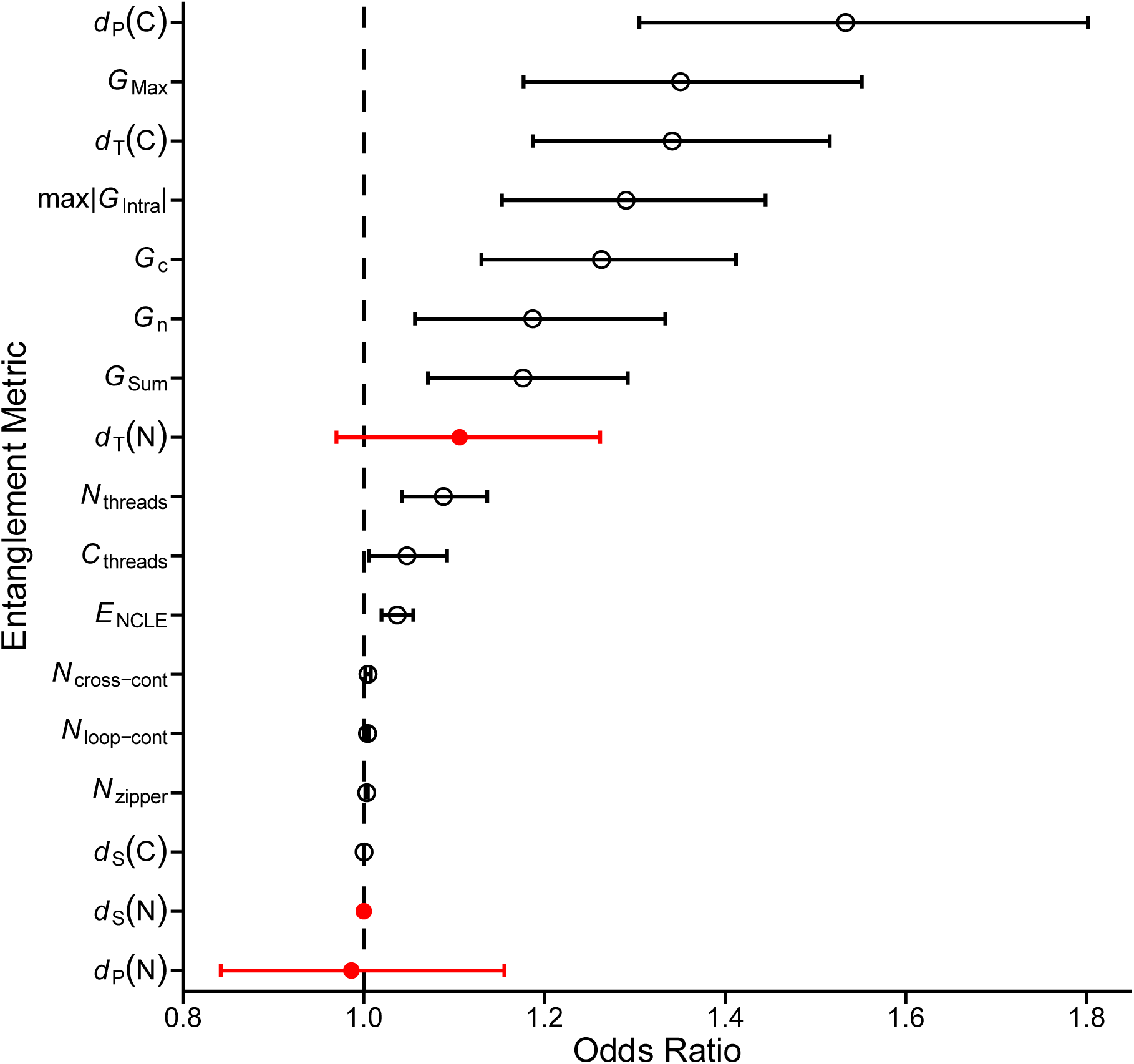
Entanglement structural metrics and disease association. Plot of odds ratios (ORs) and 95% confidence intervals from a logistic regression model (Eq. S11) measuring the association between individual entanglement structural metrics and disease status (metric definitions in Table S3; model results in Table S4). The dashed line indicates OR = 1; significant metrics (BH-adjusted *p* < 0.05) are shown as open black circles and non-significant metrics as filled red circles.

### Entangled proteins are 68% more likely to be associated with pathogenic mutations

Since entangled proteins are more likely to misfold than non-entangled proteins^24^, we hypothesized mutations could more easily cause them to misfold. Since destabilizing mutations are more likely to be pathogenic^30^, this hypothesis predicts entangled proteins will be more likely to harbor pathogenic mutations. To test this, we calculated the association between the presence of one or more entanglements in a native protein structure and the proportion of pathogenic mutations out of the set of missense mutations (Eq S12), controlling for protein length and gene essentiality (Table S5). Our dataset consists of 2,609 proteins, 38,689 pathogenic mutations, and 40,388 benign mutations (Data File 1). For missense mutations in proteins of typical length, we find that missense mutations in entangled proteins are 68% more likely to be pathogenic than mutations in non-entangled proteins (adjusted OR=1.68, p-value < 4.70 × 10^−117^, Wald test from logistic regression). Thus, entangled proteins tend to contain more pathogenic mutations because they can more easily misfold into states that significantly affect function. As a robustness test, we applied the same analysis but only to disease-associated proteins and find the association remains significant (OR = 1.48, p-value = 1.80 × 10^−58^).

### Entangled proteins are 111% more likely to be associated with pathogenic mutations in particular disease classes

We carried out the same analysis but restricted to each disease class and only disease-associated proteins. 20 out of the 23 disease classes showed a significant positive association between the proportion of pathogenic mutations and presence of entanglement, indicating that this relationship is conserved across various classes of disease-associated proteins (Table S6). The three largest associations were with Hemic and Lymphatic Diseases (C15) (OR = 2.11, Benjamini-Hochberg (BH) adjusted p-value = 3.89 × 10^−37^), Digestive System Diseases (C06) (OR = 2.02, BH-adjusted p-value = 1.08 × 10^−58^), and Respiratory Tract Diseases (C08) (OR = 1.77, BH-adjusted p-value = 2.06 × 10^−27^) (Figure 4) with percent differences of 111%, 102%, and 77%. We conclude that the excess of pathogenic mutations in entangled proteins is not driven by a single disease category but occurs across multiple classes.

**Fig. 4:**
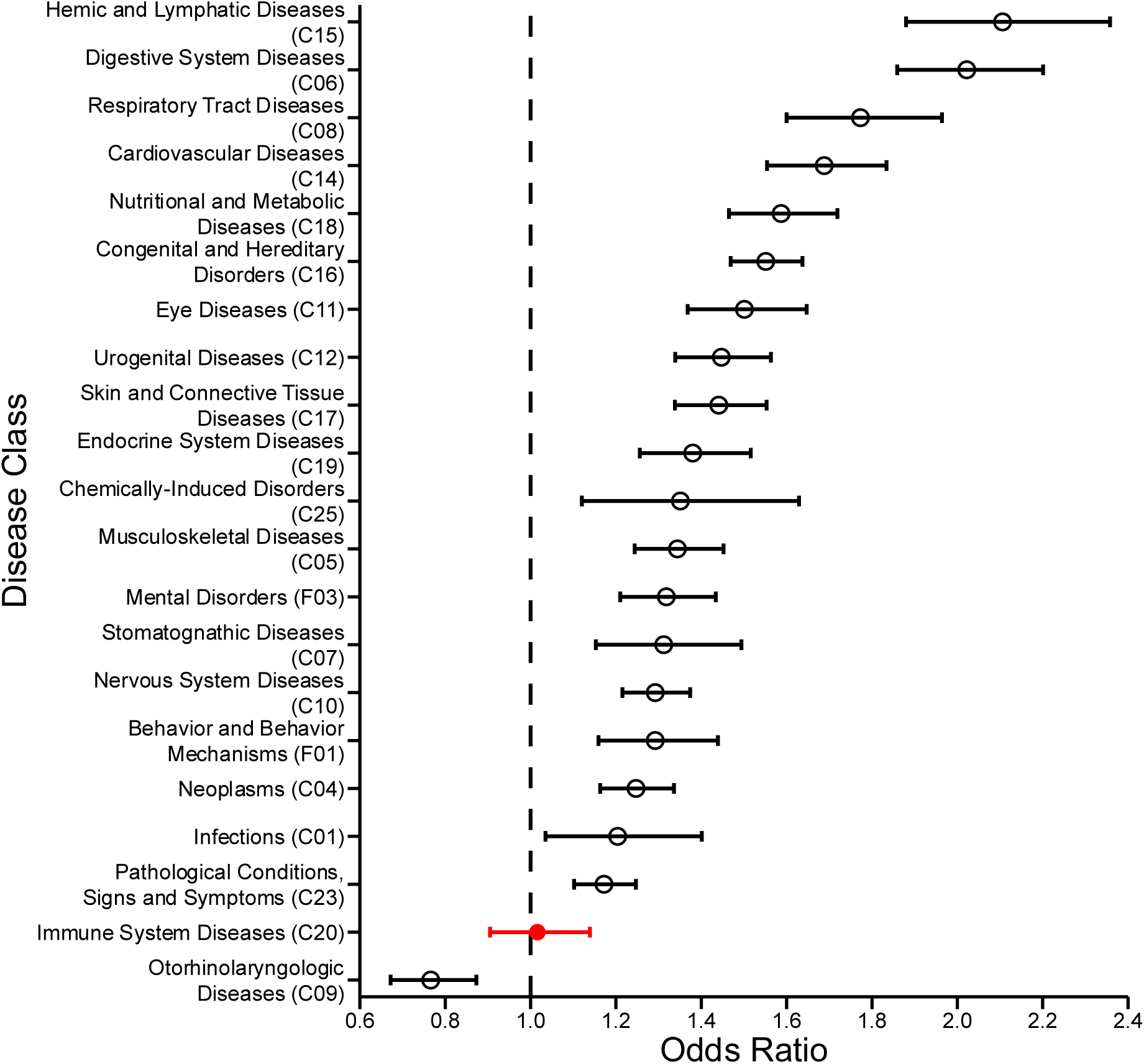
Pathogenic missense mutation enrichment in natively entangled proteins by disease class. Disease class specific enrichment of pathogenic missense mutations in entangled proteins. Points show odds ratios from logistic regression models evaluating the association between protein entanglement and the proportion of pathogenic missense mutations within each MeSH disease class, controlling for protein length and essentiality. Error bars indicate 95% confidence intervals, and the dashed horizontal line marks an odds ratio of 1 (no enrichment). Disease classes are ordered from highest to lowest odds ratio. Black and open points denote statistically significant associations after multiple testing correction, while the red closed points indicate non-significant associations. Note: Disease classes with fewer than 20 proteins were excluded from analysis. Congenital, Hereditary, and Neonatal Diseases and Abnormalities (C16) was abbreviated to Congenital and Hereditary Disorders (C16) for display purposes.

### Entangled regions along the primary structure are 64% more likely to be associated with pathogenic mutations

Simulations indicated native entanglements predominantly misfold due to premature loop closure before the threading segment is properly positioned within it^24^. We hypothesized that mutations in the entangled region of a protein are more likely to cause misfolding than non-entangled regions (Supplementary Section 1.2.3.). This hypothesis predicts that protein regions composing native entanglements are more likely to harbor pathogenic mutations as compared to non-entangled regions. To test this prediction, we calculated the association of entangled regions with pathogenic mutations; to do this we used a logistic regression model (Eq. S14) which takes as an argument whether a given residue in a protein is involved in an entanglement or not, and predicts the probability that a missense mutation in that region will be pathogenic as compared to non-pathogenic. This analysis controls for protein length and essentiality as potential confounding factors. Our dataset consists of the 2,190 unique proteins that contain one or more entanglements, 519,513 residues that are identified as taking part in an entanglement, 1,336,327 non-entangled residues, 25,502 unique residue positions with at least one pathogenic mutation, and 1,830,338 unique residues with only benign mutations or no mutations at all. We find that mutations in entangled regions have 64% higher odds of being pathogenic compared to mutations in non-entangled regions (adjusted OR = 1.64, p-value < 1.00 × 10^−300^, Wald test from logistic regression, Table S7). This result suggests that these pathogenic mutations are directly contributing to a protein’s loss of function by enhancing the likelihood of entanglement misfolding.

### Rank ordering individual proteins as potential therapeutic targets

We would like to know which proteins cause disease due to entanglement misfolding. Because observational data cannot establish causation, we use a model-based approach to prioritize proteins whose predicted disease-association probabilities differ substantially, with and without the presence of entanglements and therefore entanglement-related characteristics. That is, for an entangled protein with a set of characteristics ***Z***, we consider a counterfactual protein with the same characteristics ***Z*** but without the entanglement and quantify the difference in predicted disease risk between the two.

Specifically, we fit two random forest models: one trained on proteins with an NCLE (using NCLE structural features *U* and covariates *Z*) and one trained on proteins without an NCLE (using covariates ***Z*** only) (Supplementary Section 1.5.1). For each protein *i*, the measure of the effect size of its NCLE structural properties on disease association is 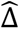 where *p̂*_1_ is the predicted disease probability for NCLE proteins (*p̂*_1_) and (*p̂*_0_) is the counterfactual probability for a non-NCLE proteins with the same other protein characteristics (*p̂*_0_). Proteins with larger 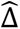 values show a larger model-predicted increase in disease-association probability when entanglements are present, suggesting a potentially stronger link to disease.

With length, mean Solvent Accessible Surface Area (SASA), mutation burden, and essentiality status being used as covariates, we report the estimated 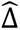 values for proteins containing native NCLES in Data File 2. The maximum estimated 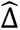 value is 0.87, reflecting a substantial increase in predicted disease association probability due to entanglement-related structural features relative to their counterfactual matched non-entangled proteins that shared similar covariates. The top scoring proteins in Data File 2, we conjecture, represent potential therapeutic targets. If true, then preventing their misfolding will result in more functional protein and the corresponding disease could be cured.

### Disease-associated entangled proteins are 2.5 times more likely to misfold

Next, we sought to understand why entangled proteins are more frequently associated with disease and harbor a greater number of pathogenic mutations. Our hypothesis is that misfolding is more likely to occur in these proteins, especially when perturbed by mutations, and lose their function. This hypothesis predicts that, compared to similar non-entangled, non-disease-associated proteins, natively entangled disease-associated proteins will exhibit higher rates of misfolding. To test this prediction, we assembled two sets of proteins for simulation. The first set consisted of 23 proteins selected from some of the top scoring proteins in Data File 2 that were disease associated and contained native NCLEs (Supplementary Section 1.5.2.). The second set was their pair-matched proteins that were not disease associated and did not contain native NCLEs (Supplementary Section 1.5.2.; Table S8). For each protein we performed coarse-grained molecular dynamics refolding simulations from a high temperature (to unfold them) and then temperature quenched them to 310 K to estimate their misfolding propensities (Supplementary Section 1.5.5). The proteins in the entangled-disease group consistently showed greater populations of misfolded states (median = 19.9%, 95% CI = [12.0%, 54.9%], bootstrap resampling of the median with 10^6^ iterations) compared to the non-entangled, non-disease proteins (median = 8.0%, 95% CI = [4.0%, 12.1%], bootstrap resampling of the median with 10^6^ iterations). The difference between these two medians is statistically significant (p-value = 2.5×10^-3^, one-sided Brunner–Munzel test^31^). Thus, the entangled, disease-associated proteins exhibit a 2.5-fold increase in misfolding propensity compared to non-entangled, non-disease proteins in this model (Fig. 5). We conclude that natively entangled, disease-associated proteins could arise from the increased propensity of these entanglements to misfold and the resulting loss of their function.

**Fig. 5:**
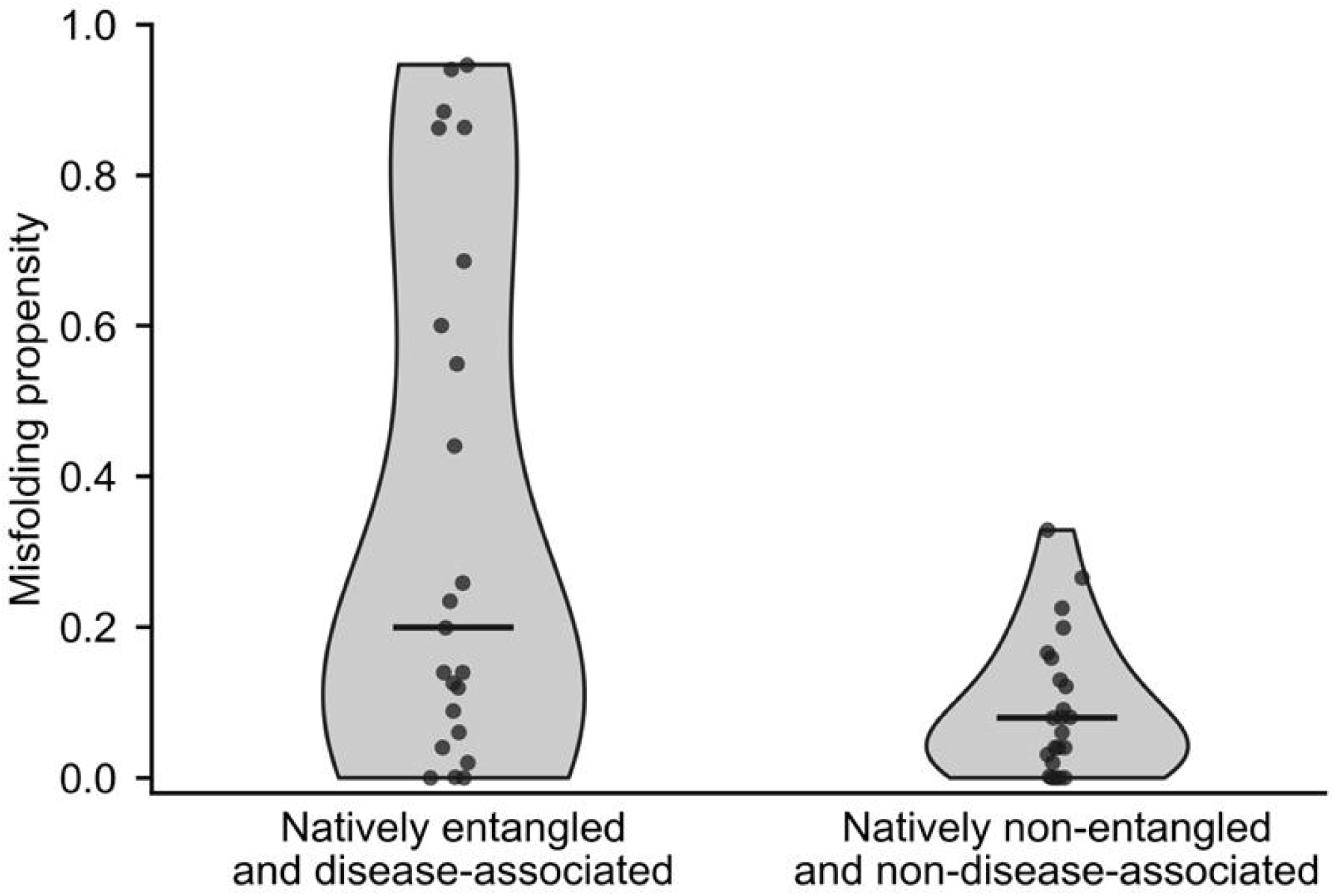
Misfolding propensity of the top scoring proteins and their control match. Misfolding propensity comparison between top 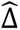 natively entangled, disease-associated proteins and their size matched non-entangled, non-disease controls (sample size n = 23 proteins per group; proteins listed in Table 8). Points represent individual proteins, horizontal bars indicate group medians, and violin shapes show the distribution of simulated misfolding proportions.

### Results are robust to parameter variation and structural data source

The DisGeNET analyses we have reported are based on using a 50^th^ percentile threshold on the disease association score. That is, for a gene to be identified as being associated with a disease its ‘S’ score, the DisGeNET-defined disease association score, must be above the median value of S across all proteins reported in DisGeNET (Supplementary Section 1.1.1.). To test the robustness of some of our key findings to variation in this threshold we reran our first three analyses using a 75^th^-percentile threshold. Our conclusions are robust. We find that entangled proteins remain positively associated with disease (OR = 1.73, p-value = 8.83 × 10^−31^) at the 75^th^ percentile (Tables S9); in the 23 disease classes, 19 now exhibit an entanglement disease association, and of these 17 are the same as those found at the 50^th^ percentile (Table S10); and in terms of entanglement features, the same 14 structural features are positively associated with disease (Table S11). Thus, the broad conclusions we report in this study are robust to moderate variation in the threshold for defining whether a gene is association with disease.

## Discussion

This study demonstrates that proteins containing native NCLE’s are more likely to be associated with disease, specific disease classes, and pathogenic mutations. Furthermore, these proteins are more likely to misfold compared to similar proteins without native NCLEs according to our simulations^24^. This suggests a new paradigm in which entanglement misfolding^24,32–34^ is a disease-causing mechanism or at least an essential step in the chain of events leading to some diseases. Previous evidence indicated that entanglement misfolding is widespread in vitro^24^. It is therefore possible that this type of misfolding is also a widespread cause of disease.

A molecular model that can explain these statistical associations are that proteins with native entanglements are inherently more prone to misfolding due to the extra complexity of folding portions of the protein backbone into a NCLE^24^. A particular series of events along the folding pathway must happen to properly fold an NCLE, otherwise the protein molecule can fall into a misfolded state^33,35^. The crucial step is the proper positioning of the thread before the loop closes – enclosing the thread and forming the native NCLE^33,35^. If the loop closes before this, then the thread is destined to remain outside the loop – misplaced and misfolded - as the timescale of threading the loop can be quite long relative to many other biological processes. Misfolded proteins are likely to compromise function, leading to inefficiencies or failure of the subcellular or cellular processes they take part in^36,37^. Thus, proteins with native NCLEs are more likely to be associated with disease and disease classes because they are more likely to misfold and malfunction. Further, pathway probabilities of different folding routes often depend sensitively on the protein sequence^35,38–40^, and the corresponding interaction energies between residues^24^. Therefore, native NCLE folding – with its required sequence of events to form - may be more easily altered by mutations than proteins lacking these structures, resulting in higher chances that a mutation will promote their misfolding. From this perspective, proteins with native NCLE’s should be enriched in pathogenic mutations.

This molecular model suggests why particular structural features of native entanglements are positively associated with disease (Table S4). For example, for each unit increase in G_max_, which describes the maximum observed complexity of an entanglement within a native structure, there is an increased likelihood that protein is associated with disease. Our molecular model says this arises because more complex entanglements are more likely to misfold, because they require a larger number of precise steps to occur. In another instance, as the number of NCLE’s per protein increases, so too does the probability that protein is associated with disease. The molecular model would suggest that as the number of native entanglements increases, the cumulative probability that one or more misfold increases, leading to a higher probability of loss of function. Taken together, these results indicate that within the set of proteins containing NCLEs, those with the largest number of complex native entanglements are most likely to be associated with disease, probably because they are the most likely to misfold and malfunction.

Our findings open a new space of therapeutic protein targets and suggest a general strategy for treating a variety of diseases. Disease-associated proteins with native entanglements and unknown mechanisms are more likely to cause disease through entanglement misfolding. To create a list of potential therapeutic targets this knowledge can be further supplemented by other sources of information, such as molecular simulations that suggest a proteins’ misfolding pathway, and structural biology data – such as limited proteolysis – that can detect such misfolding. We provide a list of therapeutic targets in Table S8 but note that it does not incorporate this additional information. Correcting the misfolding of these proteins by small molecules, under this paradigm, would restore their function and thereby treat the resulting disease. Indeed, a number of loss-of-function diseases have already been successfully treated by creating small molecules that help proteins correctly fold^41,42^.

The statistical associations between NCLEs and disease are observational and not sufficient to demonstrate causation. Such associations, however, are often a necessary condition for there to exist a causal relationship. Future experimental studies could establish causality by establishing mechanism and demonstrating that when this misfolding is corrected, disease phenotype is reduced. These represent significant efforts, but we believe they are fruitful areas of future research due to the potential widespread relevance of this type of misfolding to disease and the new therapeutic targets it suggests.

## Methods

### Data retrieval and pre-processing

Reviewed *Homo sapiens* genes were retrieved from UniProt (April 14^th^, 2022) and mapped to Entrez IDs using the UniProt ID mapping tool, retaining entries with Entrez IDs (Data File 1). Unmapped genes reflected missing RefSeq cross-references/GeneID cross-references and annotation-pipeline differences between UniProt and NCBI. Disease–gene associations were obtained from DisGeNET^26^, and disease status was binarized from DisGeNET scores using thresholds at the 50^th^ and 75^th^ percentiles {*t*_75_, *t*_50_} and setting *D*(*t*)_*i*_ = 1 if 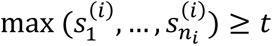 and *D*(*t*) = 0 otherwise for each threshold *t* ∈ {*t*_75_, *t*_50_}; proteins positive at *t*_50_ but negative at *t*_75_ were excluded from the *t*_75_ model to preserve a consistent reference group.

Structures were taken from AlphaFold2 (v4) and filtered by mean pLDDT, 〈*p*_LDDT_(*i*)〉 ≥ 70. SNVs were downloaded from ClinVar (GRCh38/hg38; Data File 1) with completeness filtering. Pathogenic and benign sets were defined by “ClinicalSignificance” labels without conflicting interpretations, and benign variants were enriched using v4 gnomAD PASS SNVs. ClinVar variants were mapped to UniProt canonical isoform positions via ProtVar (Data File 1) and used for downstream analyses. Gene essentiality was obtained from DepMap CRISPR gene effect datasets (downloaded October 2025) and encoded as essential/nonessential/not tested. Residue SASA was computed using Shrake–Rupley in MDTraj (probe radius 0.14 nm; 960 sphere points/atom) and converted from nm² to Å² by ×100.

### Entanglement detection

Native non-covalent lasso entanglements were identified by Gauss linking integration, requiring a loop and a threading segment piercing the loop plane. Loops were defined by native contacts between residues *i* and *j* (heavy atoms within 4.5 Å); a loop was deemed entangled if |*g*_*n*_(*i*, *j*)| ≥ 0.6 or |*g*_*c*_(*i*, *j*)| ≥ 0.6 (Eq. S3). Threading residues crossing the loop plane were validated with Topoly^43^ using chirality-consistency checks, with additional implementation details as in Rana et al. (2024).

Entanglement complexity was summarized by the maximum intrachain non-covalent lasso entanglement, defined as the largest absolute Gauss linking value max |*G*_Intra_| over all loop contacts; conformations were considered entangled when max |*G*_Intra_| ≥ 0.6. For each representative motif, “key” residues were those within ±3 of loop-closing contact residues (*i*, *j*) and the validated crossing residues; the entangled region was the union of this key set plus residues with *C*_*α*_ within 8 Å of any key residue, partitioned into crossing and loop-closing regions as in the supplemental methods.

### Entanglement Disease Association Analysis

Associations between entanglement presence and disease status were tested with logistic regression, adjusting for standardized protein length and essentiality. Generalized linear models were fitted in R^44^ separately for the two disease definitions *D*(*t*) based on thresholds *t*_50_ and *t*_75_. Odds ratios 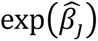 and corresponding Wald-test p-values were reported. As an unadjusted robustness check, we also performed two-sided Fisher’s exact test in R^45^. Diseases were categorized into 23 MeSH classes and class-specific logistic models were fit with the same covariates across both thresholds, applying Benjamini–Hochberg multiple testing correction across these tests.

We also assessed the association between disease and entanglement using 17 complexity-related metrics (Table S3). Each metric was first computed for individual entanglement motifs and then summarized at the protein level by taking the maximum across motifs. For each metric, we fitted a logistic regression with disease as the response variable and the standardized metric as the primary predictor, controlling for standardized length and essentiality. We used Benjamini-Hochberg procedure to obtain adjusted p-values to identify significant features.

### Entanglement Pathogenic Missense Mutation Association Analysis

The association between entanglement presence and the proportion of pathogenic missense variants was tested using a binomial regression model. The analysis was performed on proteins with at least 10 missense variants, adjusting for standardized protein length and essentiality. The same analysis was repeated on proteins that are associated with at least one disease (disease class membership defined by the *t*_50_DisGeNET threshold), and separately within each of 23 MeSH disease classes (limited to classes with ≥ 20 proteins). Benjamini–Hochberg correction was applied across the set of class-level p-values (*α* = 0.05).

A residue-level logistic regression (restricted to entangled proteins with ≥10 missense variants) was performed to study whether residues located in entangled regions are more likely to carry pathogenic missense variants. Residues were labeled as pathogenic if they carried at least one pathogenic missense variant, and as entangled if they fell within motif loop boundaries or crossing-buffer regions (as defined above).

### Coarse grained protein folding simulations

Random forest models were used to estimate a model-based disease risk contrast, defined as the difference in predicted disease-association probability with entanglement related structural features beyond other protein characteristics. Two random forest models were fitted (1,000 trees each): one on entangled proteins using entanglement features (Table S3, excluding Length) together with covariates (length, total missense count, mean SASA, essentiality), and the other on non-entangled proteins using covariates only. For each entangled protein *i*, we computed *p̂*_1*i*_, the predicted disease probability for an entangled protein based on both entanglement features covariates, and *p̂*_0*i*_, the predicted disease probability with the same covariates but without the entanglement. The estimate disease risk of protein *i* was then defined as 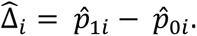

Entangled, disease-associated proteins were ranked by 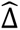 and the top 50 were matched to non-entangled, non-disease-associated controls using mean SASA, length, total missense (all standardized), and essentiality status as matching variables. Matching used the R package MatchIt^46^ (nearest-neighbor 1:1; Euclidean distance; exact matching on essentiality), and pairs were retained for simulation only if Euclidean distance <= 0.15, both proteins satisfied <20% intrinsically disordered residues (Table S8; Data File 2).

Temperature-quench folding simulations used a previously published Cα coarse-grained Gō-like model^35,47^. Each protein was unfolded at 600 K for 15 ns, cooled to 310 K, then simulated at 310 K for 50 independent 1.5 μs trajectories with a Langevin thermostat (15 fs timestep; friction 0.05 ps⁻¹), saving frames every 5,000 steps. Simulations were performed using OpenMM^48^ v8.0. Per-frame nativeness was quantified by fraction of native contacts Q and order parameter G; clustering used Kmean++ microstates^49^ (100) and PCCA+ macrostates^50^ (count chosen by visual inspection of log-probability plots) with Deeptime^51^. Misfolding propensity was computed as the fraction of non-native conformations in the final 200 ns, with a frame called misfolded only if it belonged to a non-native cluster, had Q > 0.6, and had G > 〈G〉 + 3σ (〈G〉 and σ from the native cluster), following prior criteria.

## Resource availability

## Lead contact

Requests for further information and resources should be directed to and will be fulfilled by the lead contact, Edward P. O’Brien.

## Materials availability

This study did not generate new unique reagents.

## Data and code availability

The fully reproducible version of the analysis is available at https://github.com/NCEMS/NCLE-Disease-Association-Project (DOI: 10.5281/zenodo.19373314), with all associated data deposited on CyVerse (https://de.cyverse.org/data/ds/iplant/home/shared/NCEMS/working-groups/disease-entanglement-project/NCLE-Disease-Association <u>Project%20Data%20(Reduced)/Data_Cyverse</u>). This repository contains a modified version of the analysis pipeline that uses processed derivatives of proprietary data in place of raw data from DisGeNET to comply with their user agreement. For example, rather than including the original disease association scores from DisGeNET, this version uses a binary variable indicating disease association derived from those scores. This repository reproduces all results presented in the paper.

The original version of the code, which operates on the raw data as used during the initial analysis, is available at https://github.com/NCEMS/NCLE-Disease-Association-Project-Full (DOI: 10.5281/zenodo.19373330), with partial data deposited on CyVerse (https://de.cyverse.org/data/ds/iplant/home/shared/NCEMS/working-groups/disease-entanglement-project/NCLE-Disease-Association-Project%20Data%20(Full)/Data_Cyverse). This repository is not independently executable, as certain proprietary data from DisGeNET have been removed to comply with their user agreement. Users who wish to run this version must obtain the required data directly from DisGeNET (https://www.disgenet.org/) under their own license agreement.

## Acknowledgements

This work was supported by the National Science Foundation through the National Synthesis Center for the Emergence of Molecular and Cellular Sciences NCEMS (DBI-2335029), as well as the National Institutes of Health (R35-GM124818). M.F.A. was supported by a 2025/2026 Rising Researcher Grant from the Penn State Institute for Computational and Data Sciences (RRID:SCR_025154). Computational parts of this project were carried out using the Roar Core Facility at the Penn State Institute for Computational and Data Sciences (RRID:SCR_026424).

## Contributions

M.F.A.: Investigation, Methodology, Formal analysis, Visualization, Validation, Data curation, Writing—original draft, Writing—review and editing. I.S.: Methodology, Data curation, Visualization, Writing—original draft, Writing—review and editing. Q.V.V.: Methodology, Data curation, Visualization, Writing—original draft, Writing—review and editing. P.T.: Data curation, Writing—review and editing. J.D.S.: Data curation, Writing—original draft, Writing—review and editing. H.S.: Conceptualization, Formal analysis, Investigation, Methodology, Project administration, Resources, Supervision, Validation, Writing—original draft, Writing—review and editing. E.O.: Conceptualization, Formal analysis, Funding acquisition, Investigation, Methodology, Project administration, Resources, Supervision, Validation, Writing—original draft, Writing—review and editing.

## Declaration of interest

The authors declare no competing interests.

## Declaration of generative AI and AI-assisted technologies in the manuscript preparation process

During the preparation of this work the authors used ChatGPT for proof-reading and editing. After using this tool, the authors reviewed and edited the content as needed and take full responsibility for the content of the published article.

## Notes

https://github.com/NCEMS/NCLE-Disease-Association-Project

https://de.cyverse.org/data/ds/iplant/home/shared/NCEMS/working-groups/disease-entanglement-project/NCLE-Disease-Association-Project%20Data%20(Reduced)/Data_Cyverse

https://github.com/NCEMS/NCLE-Disease-Association-Project-Full

https://de.cyverse.org/data/ds/iplant/home/shared/NCEMS/working-groups/disease-entanglement-project/NCLE-Disease-Association-Project%20Data%20(Full)/Data_Cyverse

### Competing Interest Statement

The authors have declared no competing interest.

